# Prevailing role of mucosal immunoglobulins and B cells in teleost skin immune responses to bacterial infection

**DOI:** 10.1101/2020.09.21.305920

**Authors:** Xiao-Ting Zhang, Yong-Yao Yu, Hao-Yue Xu, Zhen-Yu Huang, Xia Liu, Jia-Feng Cao, Kai-Feng Meng, Zheng-Ben Wu, Guang-Kun Han, Meng-Ting Zhan, Li-Guo Ding, Wei-Guang Kong, Nan Li, Fumio Takizawa, Sunyer J Oriol, Zhen Xu

**Affiliations:** Department of Aquatic Animal Medicine, College of Fisheries, Huazhong Agricultural University, Wuhan, Hubei, 430070, China; State Key Laboratory of Freshwater Ecology and Biotechnology, Institute of Hydrobiology, Chinese Academy of Sciences, Wuhan, Hubei, 430072, China; Faculty of Marine Science and Technology, Fukui Prefectural University, Obama, Fukui 917-0003, Japan; Department of Pathobiology, School of Veterinary Medicine, University of Pennsylvania, Philadelphia, Pennsylvania, 19116, USA; Laboratory for Marine Biology and Biotechnology, Qingdao National Laboratory for Marine Science and Technology, Qingdao, Shandong, 266071, China

## Abstract

The skin of vertebrates is the outermost organ of the body and serves as the first line of defense against external aggressions. In contrast to mammalian skin, that of teleost fish lacks keratinization and has evolved to operate as a mucosal surface containing a skin-associated lymphoid tissue (SALT). Thus far, IgT representing the prevalent immunoglobulin (Ig) in SALT have only been reported upon infection with a parasite. However, very little is known about the types of B cells and Igs responding to bacterial infection in the teleost skin mucosa, as well as the inductive or effector role of the SALT in such responses. To address these questions, here we analyzed the immune response of trout skin upon infection with one of the most widespread fish skin bacterial pathogens, *Flavobacterium columnare*. This pathogen induced strong skin innate immune and inflammatory responses at the initial phases of infection. More critically, we found that the skin mucus of fish having survived the infection contained significant IgT-but not IgM- or IgD-specific titers against the bacteria. Moreover, we **d**emonstrate the local proliferation and production of IgT^+^ B-cells and specific IgT titers respectively within the SALT upon bacterial infection. Thus, our findings represent the first demonstration that IgT is the main Ig isotype induced by the skin mucosa upon bacterial infection, and that because of the large surface of the skin, its SALT probably represents a prominent IgT inductive site in fish.

## Introduction

In contrast to five major Ig classes (IgM, IgG, IgA, IgD, and IgE) in mammals, only three Ig isotypes are present in teleosts, including IgM, IgT/IgZ, and IgD (1). Teleost IgM is the most abundant Ig class that involved in systemic immunity upon infection or vaccination (2–5). Secreted IgD (sIgD) has been identified and found coating a significant portion of the fish microbiota (6–8), however, its immune function against pathogens remains largely unknown. Teleost IgT has been demonstrated to play a specialized role in mucosal immunity akin to that of mammalian secretory IgA (sIgA) (6–10). Similar to sIgA, sIgT plays a key function both in protecting fish mucosal sites against pathogens (7–12), as well as in promoting microbiota homeostasis at mucosal surfaces (13). With regards to its role in pathogen control, we and others have shown that sIgT is the main Ig induced in several fish mucosal associated lymphoid tissues (MALTs) upon pathogen insult (7–12). Thus far, pathogen-specific IgT responses have been documented in gut, skin, gill, nasal, buccal and pharyngeal mucosal areas (7–12). However, in-depth studies assessing the local induction of IgT have only been performed in some of the fish MALTs, including those of the gills, nose and buccal and pharyngeal tissues (7, 8, 11, 12). Moreover, such local responses have been assessed with the use of only one pathogen type, the parasite *Ichthyophthirius multifiliis* (Ich), a pathogen that is responsible for important losses to both wild and farmed populations of salmonid species.

At this point, nothing is known about the local induction of IgT responses in the skin of fish, by far the largest mucosal surface in these species. Moreover, pathogen-mediated IgT responses in that surface are restricted to one study that utilized the parasite Ich as the pathogen model (9), while it is unknown at this point whether bacterial pathogens are capable of inducing similar responses. Anatomically, the skin presents a similar structure in all vertebrates, which is generally composed of two layers (epidermis and dermis) (9). However, it should be noted that vertebrates of different taxonomic statuses have faced unique evolutionary pressures that have continuously shaped their own skin structures and components. These histological and componential differences in the skin of vertebrates are reflected also in their specific immune strategies (14). It is worth noting that the term SALT was first used in reference to mammals (15) and that in these species, the main cellular constituents of the skin epidermis are specialized epithelial cells known as keratinocytes, Langerhans cells, and T cells (16). Moreover, the skin dermis of mammals is home to a diverse array of specialized immune cells, including antigen-presenting dermal dendritic cells (DCs), T cells, B cells, and natural killer (NK) cells as well as mast cells, monocytes, and macrophages (17), which indicates that many immunological processes taking place in the dermis layer of the mammalian SALT (9). Interestingly, teleost fish have also been shown to contain a SALT (2). However, in contrast to mammalian skin, that of teleosts behaves as a mucosal surface since it lacks keratinization, harbors abundant mucus, and its living epithelial cells are in direct contact with the water environment where they live. While the fish dermis is mainly composed of dense connective tissue with a large amount of collagen fibers and is devoid for the most part of immune cells, its epidermis has been shown thus far to contain B cells, T cells, macrophages and granulocytes as the main immune cells (9). Moreover, it is in the epidermis where significant immune responses have also been reported (18, 19). IgT responses in the skin were first described by us, where we showed that IgT-specific responses against a parasite were chiefly detected in the skin mucus, whereas IgM-specific titers were for the most part observed in the serum (9). Whether IgT and IgT^+^ B cell responses are locally induced in the skin is an important question that remains to be addressed. In that regard, it is important to note that in mammals not all IgA responses are induced locally in all existing mucosal sites (20). For example, IgA found in some mucosal areas (e.g. female genital tract) and secretory glands (e.g. salivary glands and lacrimal glands) is derived from B cells that are locally induced in other mucosal sites (e.g. nasopharynx-associated lymphoid tissue (NALT) and gut-associated lymphoid tissue (GALT)) (21–23).

As stated above, only one report has described the induction of skin IgT-specific titers against a pathogen (Ich), and local IgT responses have only been assessed, with the use of the same pathogen, only in some fish mucosal sites, but not the skin. Prior to the discovery of the role of IgT in mucosal immunity, several studies in teleosts showed the induction of modest IgM-specific titers against several bacterial pathogens. More recently, a couple of studies were unable to detect IgM-specific titers in the skin mucus of rainbow trout exposed to *Flavobacterium pschrophilum* which prompted the authors to suggest that IgT might play a role against this bacterial pathogen. Thus, in this study we aimed to evaluate the specific contributions of the different fish Igs to skin bacterial pathogen in a representative teleost, and to assess whether such responses could be induced locally in the skin-associated lymphoid tissue (SALT). Here we choose *Flavobacterium columnare* (*F. columnare*), one of the most widespread fish skin bacterial pathogens. *F. columnare* is a long Gram-negative rod in the family *Flavobacteriaceae*, one of the main phyletic lines within the Bacteroidetes group from the domain Bacteria (24), which can survival in both fresh and brackish water throughout the world (25).This pathogen can cause columnaris disease that affects a variety of freshwater fishes, leading significant losses both in wild and farmed fish populations (26–32). Columnaris disease is characterized by a pronounced erosion and necrosis of mucosal tissues including the fin, skin and gills and it can cause particularly high mortalities. Secreted enzymes, such as proteases and chondroitin sulfate lyases, have been potential *F. columnare* virulence factors, contributing to the branchial and cutaneous necrosis (33, 34). While we have shown in the past that *F. columnare* induces dominant IgT- and IgM-specific responses in the gill mucosa and serum of rainbow trout respectively, thus far, little is known with regards to the pathogenesis of columnaris in the fish skin as well as the innate and adaptive immune mechanisms induced by this pathogen in that surface upon infection.

To assess the mucosal immune responses induced in the skin by *F. columnare*, we infected rainbow trout with a newly constructed fluorescent version of this pathogen following a bath exposure strategy (35). Our findings show that trout skin elicited strong innate and adaptive immune responses to *F. columnare*. More critically, we show that *F. columnare*-specific Ig responses in the skin mucus were mainly represented by the IgT isotype and that such responses were induced locally as shown by the local proliferation and accumulation of IgT^+^ B cells. Moreover, when cultured in vitro, skin explants produced *F. columnare*-specific IgT titers. These data show for the first time that the use of a fluorescent version of *F. columnare* which was critical to further understand the pathogenesis of this important fish pathogen. More importantly, we demonstrate the previously undiscovered capacity of the fish skin of inducing potent mucosal IgT-specific responses against a bacterial pathogen and that such responses are induced locally upon mucosal infection.

## Materials and methods

### Ethics statement

All experimental protocols were performed in accordance with the recommendations in the Guide for the Care and Use of Laboratory Animals of the Ministry of Science and Technology of China. They were approved by the Scientific Committee of Huazhong Agricultural University (permit number HZAUFI-2017-013). All efforts were made to minimize the suffering of the animals.

### Fish

Rainbow trout (∼ 5 g) were obtained from a fish farm in Shiyan (Hubei, China) and were maintained in aerated aquarium tanks with a water recirculation system including internal biofilters and thermostatic temperature control. Fish were acclimatized for at least 2 weeks at 16°C, and fed daily with commercial trout pellets at a rate of 1% biomass during the whole experiment periods. The feeding was terminated 2 days prior to sacrifice.

### F. columnare *culture and construction of GFP-F*. columnare *G_4_ strain*

The strain G_4_ of *F. columnare* used in this study was provided by Professor Nie (36) and conserved at -80 °C. *F. columnare* strains were grown at 28 °C in Shieh medium or on Shieh agar. The GFP-*F. columnare* G_4_ strain was constructed by using the method previously described by Li et al (36). Briefly, a fused fragment carrying P*ompA* of *F. johnsoniae* (*Flavobacterium johnsoniae*) and *gfp* was amplified by PCR using the primers pr44-Bam (CGCGGATCCGGCAGCGCATACCAAAGAACACTTAG) and pr38-Pst (GCTAGCTGCAGCAGATCTATTTGTATAGTTCATCCA) with pAS43 as template. The PCR product was digested with BamHI and PstI and cloned into the corresponding sites of pCP23 to generate plasmid pCP-gfp. This plasmid was transferred into *Escherichia coli* (*E. coli*) S17-1 λ pir and then was introduce into G_4_ by conjugation. The GFP-labeled strain was screened on Shieh agar containing 1 μg/ml tobramycin and 10 μg/ml tetracycline.

### Growth curve determination

To compare the growth rates of GFP-*F. columnare* and wild-type *F. columnare* G_4_ strain *in vitro*, both strains were streaked on Shieh agar at 28 °C for 24 h. The colonies were then inoculated into Shieh broth and incubated at 28 °C with shaking at 150 rpm. The optical density was measured at 600 nm with 2ml cuvette using NanoPhotometer^®^ NP80 (Implen, Germany). Until the OD_600_ reached 0.4, 500 μl of such early-exponential-phase were inoculated in 25 ml Shieh broth and incubated at 28 °C with shaking at 150 rpm. The OD_600_ of culture liquid from each group was measured every 1 h for 13 h.

### Rainbow trout challenges

The method used for *F. columnare* infection were described previously (19) with slight modification. Fish (∼ 5 g per fish) were randomly chosen and exposed by immerse with wild-type or GFP-*F. columnare* at a final concentration of 1×10^6^ CFU/ml for 4 h at 16 °C, and then migrated into the aquarium containing new aquatic water. To detect the localization of *F. columnare* in trout, 30 fish were infected with GFP-*F. columnare*, and tissue samples including skin, gill and fin were collected at 1, 2, 4, 7 and 14 dpi. For the study of immune responses in trout, there were two types of challenges were performed. In the first group (infected fish), 60 fish were challenged by wild-type G_4_ strain. Tissue samples including skin, head kidney, and spleen were collected at 1, 2, 4, 7, 14, and 28 dpi. Skin mucus and serum were taken 4 weeks later as infected group (fish survived 28 days after one exposure). In the second group, another 60 fish were monthly exposed to wild-type *F. columnare* for 75 days with the final concentration of 1×10^6^ CFU/ml for 4 h at 16 °C. Fish samples mentioned above were taken 2 weeks after the last challenge as survivor group (fish survived 75 days after three exposure). Both experiments were performed at least three independent times. As a control group (mock infected), the same number of fish were maintained in similar tanks and were exposed to the same culture medium without pathogens.

### *The detection of* F. columnare *in trout*

To detect the *F. columnare* in rainbow trout after challenge, different tissue samples including skin, gills and fins were collected, weighed and homogenized in tissue lysis buffer using steel beads and shaking (60 HZ for 1 min) by TissueLyser II (Jingxin Technology). Then the DNA of different tissue samples was extracted and amplified by PCR using the 16S rRNA specific primers of *F. columnare*. The PCR products were extracted from a 2% agarose gel, and images were acquired using a Gel Doc XR^+^ (Bio-Rad, USA). Moreover, skin tissue samples were collected from both control and infected fish and homogenized in 1 ml PBS (filtered with 0.22 μm) for coating plates, and the colonies were observed using a fluorescence microscope.

### RNA isolation and quantitative real-time PCR (qRT-PCR) analysis

Total RNA was isolated by TRIZol (Invitrogen) method as we previously described (37). Approximately 2 μl of RNA (1000 ng in total) was reverse transcribed into cDNA with Hifair III 1st Strand cDNA Synthesis SuperMix for qPCR (YEASEN) following the manufacturer’s instructions. And then, qRT-PCR was performed by with SYBR green qPCR mix (Monad) at 95°C for 5 min, and 41 cycles of 95°C 10 s, 58°C 30 s. We calculated the relative fold changes by using the 2^-ΔΔCt^ method and EF-1α was used as an internal control. The specific primers used for this experiment are list in Supplemental Table I.

### RNA-Seq libraries and differential expression analysis

Nine samples from rainbow trout was used for the RNA-Seq libraries construction and analyzed as we previous described (38). RNA-Seq data generated by Illumina paired-end sequencing was filtered and only reads mapped to one site was used for further analysis. Genes with FDR ≤ 0.05 and |log2 (fold-change) | ≥ 1 were considered as the differentially expressed genes (DEGs).

### Collection of serum and skin mucus

MS-222 was used to anesthetize the trout, and after that, the serum was obtained by ultracentrifugation (10). As for the skin mucus, we used the method described by Xu et al (9). Briefly, we transferred the scraping from the surface of skin into an Eppendorf tube, and then blew it repeatedly through the injector. Subsequently, the skin mucus was collected by centrifuge. To remove the skin bacteria, we centrifuged the previously treated skin mucus by 10,000 *g* ultracentrifugation and then the skin mucus supernatant was obtained.

### Western blot

Skin mucus and serum under non-reducing condition were separated by 4-15% SDS-PAGE Ready Gel (Bio-Rad) according to previous research (7). The samples were detected by western blot analysis using primary anti-trout IgT (rabbit polyclonal Ab (pAb)), anti-trout IgM (mouse monoclonal antibody (mAb)) or biotinylated anti-trout IgD (mouse mAb), respectively. The binding of primary antibody was detected by secondary antibodies and observed by GE Amersham Imager 600 Imaging System (GE Healthcare).

### Histology, light microscopy and immunofluorescence microscopy studies

The paraffin sections of tissues were obtained for histology or immunofluorescence study as previous report (38). To detect the histological structure and mucus cells in skin, the sections were stained with H&E and AB. To observe the distribution of GFP-*F. columnare* in skin, gills and fins, the sections were stained with DAPI (4′, 6-diamidino-2-phenylindole; 1 μg/ml; Invitrogen). In the immunofluorescence study, polyclonal rabbit anti-trout IgT (0.5 μg/ml), monoclonal mouse anti-trout IgM (1 μg/ml), or polyclonal rabbit anti-trout pIgR antibody (0.5 μg/ml) were used to assess the IgT^+^ and IgM^+^ B cells or pIgR^+^ cells in skin sections as previous study (9). All images were gained by Olympus BX53 fluorescence microscope (Olympus), and processed by iVision-Mac scientific imaging processing software (Olympus).

### Proliferation of B cells in the skin of trout

The proliferation of B cells in skin and head kidney were assessed in control and survivor fish by injection wih EdU (Invitrogen) as previously reported (7). 24 hours later, the leucocytes from trout skin and head kidney were isolated for flow cytometry study using the existing methodology as described previously (7 – 9). To detect IgT^+^ or IgM^+^ B cells, approximately 5×10^6^ cells were incubated with anti-trout IgT (IgG 2b, 1 μg/ml) or mouse anti-trout IgM (IgG1, 1ug/ml), and the secondary antibodies were incubated subsequently. The EdU^+^ cells were then stained according to the manufacturer’s instructions (Click-iT EdU Alexa Fluor 647 Flow Cytometry Assay Kit, Invitrogen). Using a CytoFLEX flow cytometer (Beckman coulter) and FlowJo software (Tree Star), the percentages of EdU^+^ cells within the IgM^+^ or IgT^+^ B cells populations were presented.

### Tissue explants culture

Firstly, fish were treated with MS-222 for conveniently removing blood from the caudal vein. Then, the tissues of head kidney, spleen and skin (about 50 mg) were taken from each fish, and soaked with 75% ethanol 15 s to kill the surface bacteria. Thereafter, the tissues were submerged twice in PBS and moved to 400 μl prepared DMEM medium (Invitrogen). Lastly, tissues in the medium were cultured (5% CO_2_, 17°C) for 7 days, and the tissue culture fluid was collected and stored until use.

### *Binding of trout immunoglobulins to* F. columnare

Here, we measured the titers of *F. columnare*-specific Igs in fluid samples (skin mucus, serum and tissue explant medium) by pull-down assay as previously described (7). Briefly, bacteria were incubated with fluid samples for 4 h at 4 °C and washed three times with PBS. Then the bound proteins were eluted with 2 × Laemmli Sample Buffer (Bio-Rad), and then analyzed by western blot as clarified above.

### Statistical analysis

An unpaired Student’s *t*-test (Excel version 11.0; Microsoft) and nonparametric Mann–Whitney *t*-test (Prism version 6.0; GraphPad) were used for analysis of differences between two groups. *P* values of 0.05 or less were considered statistically significant.

## Results

### Pathological changes in trout skin after bacterial infection

To evaluate morphological changes in trout skin, we used a bath challenge model with *F. columnare*, which results in the infection of fish mucosal tissues, and elicits strong immune responses (39). However, due to lack of specific antibody to this bacterial strain, a recombinant *F. columnare* strain containing a plasmid expressing green fluorescent protein (GFP) was constructed to gain insight into the pathogenesis and progression of the bacterial infection in fish skin (Supplemental Fig. 1A). Under the same culture conditions, the GFP-expressing and wild-type *F. columnare* strains grew at similar rate in Shieh medium (Supplemental Fig. 1B). Critically, both the GFP-expressing and wild-type *F. columnare* strains elicited very similar mortality rates when used at the same concentration (Supplemental Fig. 1C). Together, these results indicated that the growth and virulence of the GFP-expressing *F. columnare* (GFP-*F. columnare*) were not influenced by GFP or the plasmid. At 4 days post infection (dpi) with GFP-*F. columnare*, we observed that the classical symptoms of skin lessions, fin rot and gill necrosis appeared in the trout (Fig. 1A). Importantly, high expression of *F. columnare-*specific 16S rRNA accumulated in the skin, gills, and fins of trout at 4 dpi (Fig. 1B). Notably, by using fluorescence microscope, a large number of *F. columnare* with green fluorescence was observed in the fin, skin and gill tissues from 4-day infected fish, when compared to that of control fish (Fig. 1C). In addition, the tissue homogenates of trout skin from control fish and 4-day infected fish were cultured on Shieh agar, and the yellow-green color bacterial colonies showed characteristic rhizoid shapes were detected only from homogenate samples of infected trout skin (red arrows). Moreover, the single colonies isolated from the plate were grown in pure culture, and displayed characteristic elongated rod-shaped *F. columnare* bacteria with green fluorescence (Fig. 1D). By fluorescence microscopy, a time series observation of GFP-*F. columnare* in skin tissue showed that much of the bacteria was found in the skin epithelium, and that bacteria accumulated in trout skin as early as 1 dpi (Fig. 1E). Moreover, similar to these results, qRT-PCR analysis showed that the *F. columnare* load in trout skin increased significantly from 1 dpi and peaked at days 4 (Fig. 1F), which further confirmed that the bacteria succeeded in invading the skin mucosal tissue.

**FIGURE 1.**
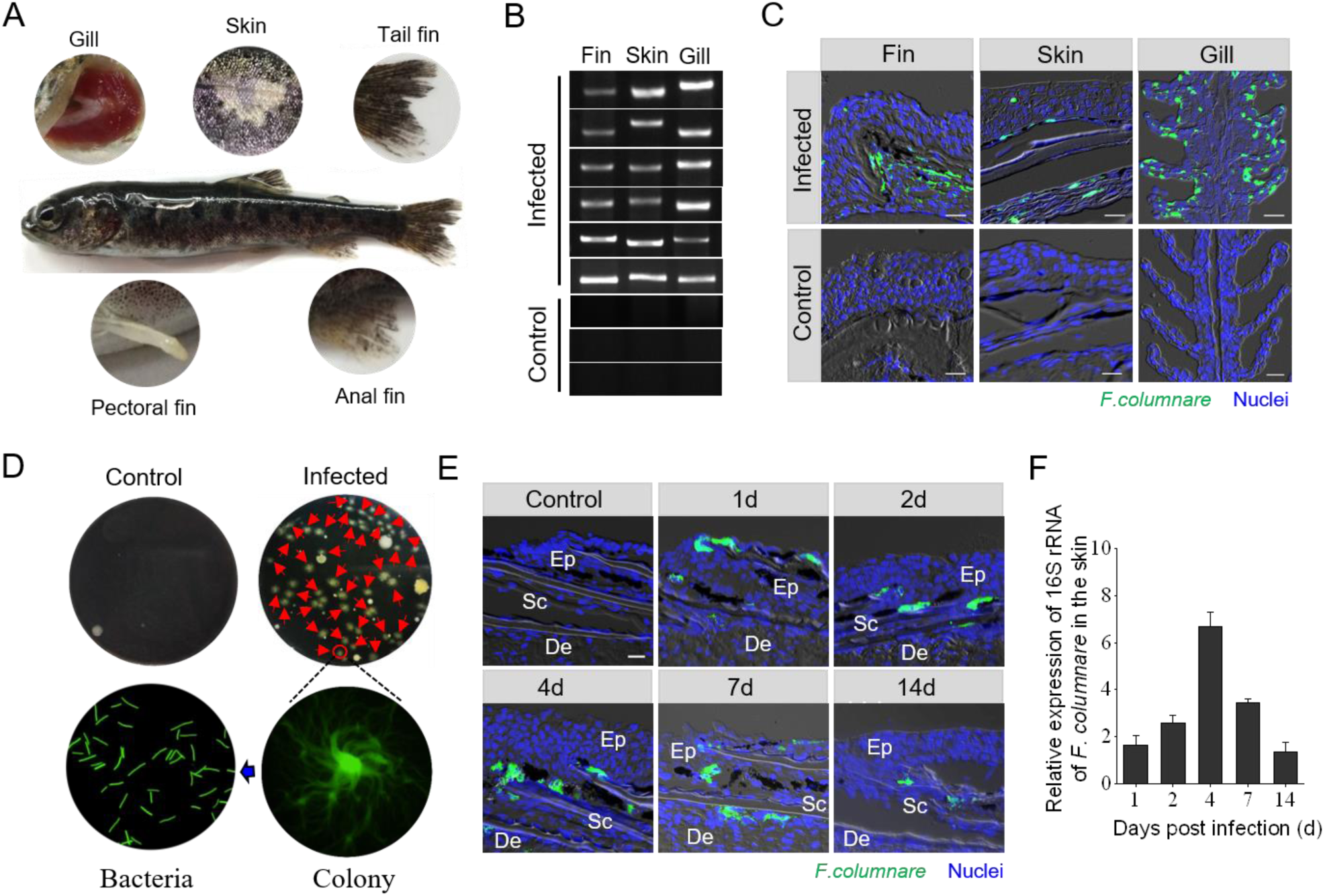
Invasion of *F. columnare* to the skin of rainbow trout. (**A**) The severe phenotype of rainbow trout was observed at 4 dpi with GFP-*F. columnare.* (**B**) The PCR products of *F. columnare* 16S rRNA gene of the fins, skin and gills tissues from three control and six infected fish at 4 dpi, were electrophoresed in 2% agarose and the bands were recorded using the gel imaging system. (**C**) Histological examination of GFP-*F. columnare* by fluorescence in trout fins, skin and gill paraffinic sections from infected fish at 4 dpi. Differential interference contrast (DIC) images show merged staining with *F. columnare* (green) and nuclei (blue). Scale bars, 20 µm. (**D**) The culture plates from trout skin of control fish (left, upper) and 4-day infected fish (right, upper), respectively. Red triangles indicate single colony of GFP-*F. columnare*. Colony image: a magnified view of circled colony from infected fish by fluorescence microscope (right, lower). Bacteria image: the observation of bacterial solution obtained by circled colony expansion (left, lower). (**E**) Localization of GFP-*F. cloumnare* in trout skin of control fish and infected fish at 1, 2, 4, 7, and 14 dpi. Ep: epidermis, De: dermis, Sc: scale. Scale bars, 20 µm. (**F**) The relative expression of 16S rRNA of *F. columnare* in infected fish versus control fish measured at days 1, 2, 4, 7, and 14 post-infection in skin of rainbow trout (*n* = 6). Data are representative of three independent experiments (mean ± SEM).

In combination with paraffin sections and staining (hematoxylin & eosin or alcian blue, H & E / AB), morphological changes were observed in the skin epidermis. We observed a conspicuous decrease in both the thickness of the skin epidermis and the number of mucus cells in the infected trout skin (Fig. 2A). Moreover, the thickness of the trout skin epidermis reduced significantly at 2 dpi, and it remained relatively stable for up to 14 days (Fig. 2B). In addition, the significantly reduced number of mucus cells in the skin epidermis occurred at 1, 2, and 7 dpi (Fig. 2C).

**FIGURE 2.**
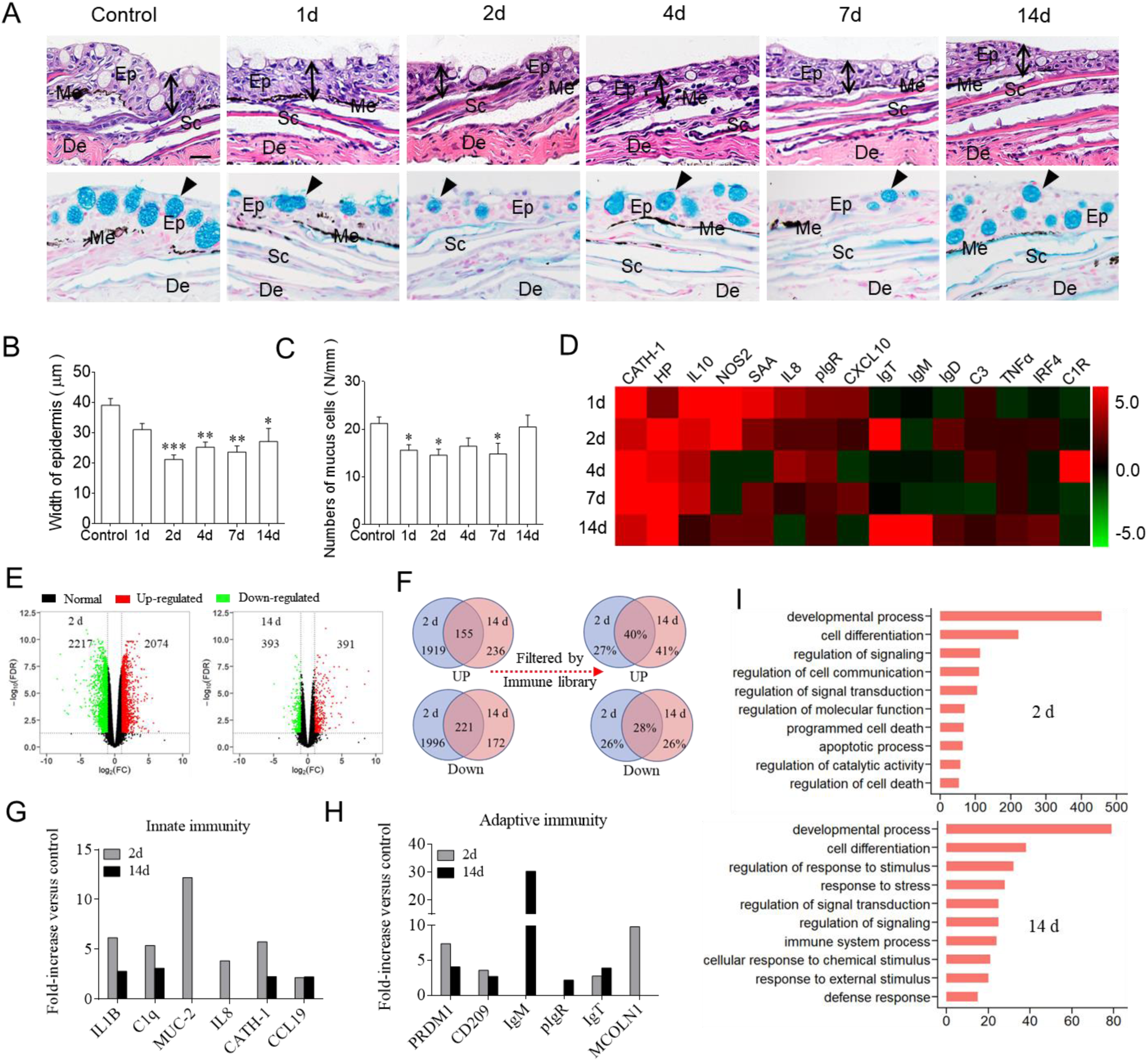
Pathological changes and immune responses in the skin of rainbow trout following *F. columnare* infection. (**A**) Histological examination by H & E and AB staining of skin from trout infected with *F. columnare* at days 1, 2, 4, 7, and 14, as well as uninfected control fish (*n* = 6). The black lines with double arrows and black triangle indicate the width of skin epidermis and the mucus secreting cells, respectively. Ep: epidermis, De: dermis, Sc: scale, Me: melanophores. Scale bars, 20 µm. (**B**) The width of skin epidermis of control and bacterial infected fish counted from A (upper) (*n* = 6). (**C**) The number of mucus cells per millimeter in the epidermis of trout skin from control and bacterial infected fish counted from A (lower) (*n* = 6). (**D**) Heat map illustrates results from qRT-PCR of mRNA expression levels for selected immune markers in *F. columnare*-infected fish versus control fish measured at 1, 2, 4, 7, and 14 dpi in the skin (*n* = 6). Color value: fold change. (**E**) Volcano plot showed the overlap of genes upregulated or downregulated in the skin of rainbow trout at days 2 (left) and 14 (right) infected with *F. columnare* versus control fish. Red spots: expression fold change of > 2 and FDR of < 0.05; Green bots: expression fold change of < 2 and FDR of < 0.05; Black spots: no difference in expression. (**F**) Venn diagrams showed the statistical results of DEGs including up-regulated and down-regulated genes and the percentage of immune genes after the DEGs filtered by rainbow trout immune genes libraries in the trout skin at days 2 and 14 infected with *F. columnare* versus control fish. (**G** and **H**) Representative innate (G) and adaptive (H) immune genes modulated by *F. columnare* infection at 2 and 14 dpi (*n* = 9). (**I**) Biological processes that were significantly altered of rainbow trout at days 2 and 14 infected with *F. columnare* versus control fish revealed by RNA-Seq studies. Fold change differences between control and bacterial infected samples were calculated using cutoff of two-fold. \**P* < 0.05, \*\**P* < 0.01, \*\*\**P* < 0.001 (unpaired Student’s *t*-test). Data are representative of three independent experiments (mean ± SEM).

### Bacterial infection elicits strong immune responses in trout skin

Using qRT-PCR, we detected the mRNA expression levels of 15 immune-related genes and cell markers, including cytokines (interleukin 8, IL8; interleukin 10, IL10; tumor necrosis factor α, TNFα; and interferon regulatory factor 4, IRF4), cathelicidin (CATH1), complement 3 (C3), poly Ig receptor (pIgR), and Ig heavy chain genes (IgT, IgM, and IgD) (Fig. 2D; primers were shown in Supplemental. Table I). These studies indicated a strong immune response occurred in trout skin (Fig. 2D), spleen (Supplemental Fig. 2A), and head kidney (Supplemental Fig. 2B) after challenge with *F. columnare*. Interestingly, significantly upregulated immune-related genes (e.g., IL10; inducible nitric oxide synthase, NOS2; serum amyloid A, SAA) were detected in trout skin as early as 1 day, while in trout head kidney (Supplemental Fig. 2A) and spleen (Supplemental Fig. 2B), immune responses were delayed upon challenge with *F. columnare*. It’s worth noting that day 2 and day 14 were most correlated in terms of the strength of the skin immune responses. Taken the time point of expression of the immune-related genes and significant histological responses from 2 dpi to 14 dpi, we chose these two time points for subsequent high-throughput transcriptome sequencing analysis. RNA-Seq libraries were constructed from nine samples that separately represented three groups (Control, 2d, and 14d) which were sequenced on an Illumina platform. Upon *F. columnare* challenge, we found significantly modified mRNA expression of 4291 genes (2074 genes upregulated and 2217 genes downregulated) at 2 dpi, and 784 genes (391 genes upregulated and 393 genes downredulated) at 14 dpi, respectively (Fig. 2E). More than 25% of DEGs indicated immune-related genes after filtering by the immune gene library of *Oncorhynchus mykiss* (*O. mykiss*) (Fig. 2F). Importantly, among the immune-related genes, we found significantly modified mRNA expression of innate immune genes (Fig. 2G; e.g. interleukins, chemokines, antibacterial peptides, and mucins) and adaptive immune genes (Fig. 2H; e.g. genes associated with antigen processing and presentation, and Igs) at both days 2 and 14. Gene ontology (GO) analysis showed that these DEGs were enriched for genes involved in developmental processes, cell differentiation, signaling transduction, programmed cell death, and immune response (Fig. 2I).These data indicated that *F. columnare* invasion could induce intense and long-term innate as well as adaptive immune responses. In order to validate the DEGs identified by RNA-Seq, we chose 12 upregulated and 12 downregulated genes for qRT-PCR confirmation. The result showed a significant correlation of the expression values determined by RNA-Seq and qRT-PCR at each time point (Supplemental Fig. 2), suggesting that the RNA-seq results performed the same accuracy as that of qRT-PCR in the determination of gene expression in *vivo*.

### Pathogen load decreased in fish skin after re-infection

Histopathology of the skin of the both infected and survivor groups showed no significant changes in the epidermis compared to the control group (Fig. 3A, 3B), and the number of mucus secreting cells also recover to normal levels (Fig. 3A, 3C). We then challenged the infected (28 dpi) and survivor fish as well as uninfected control fish by bath with 5×10^6^ CFU/ml of GFP-*F. columnare* for 4 h at 16 °C. All fish were euthanized at 4 days post challenge to evaluate bacterial load in the skin tissues from the three fish groups. Using fluorescence microscopy and qRT-PCR, we detected that upon re-infection, the infected and survivor groups had markedly limited green fluorescence signal and lower 16S rRNA expression of *F. columnare* compared with those of the control-challenged fish (Fig. 3D, 3E). These data strongly suggest that pre-exposure to *F. columnare* protects trout skin from the bacterial re-infection, suggesting that mucosal adaptive immunity had been induced in trout skin to fight pathogen invasion.

**FIGURE 3.**
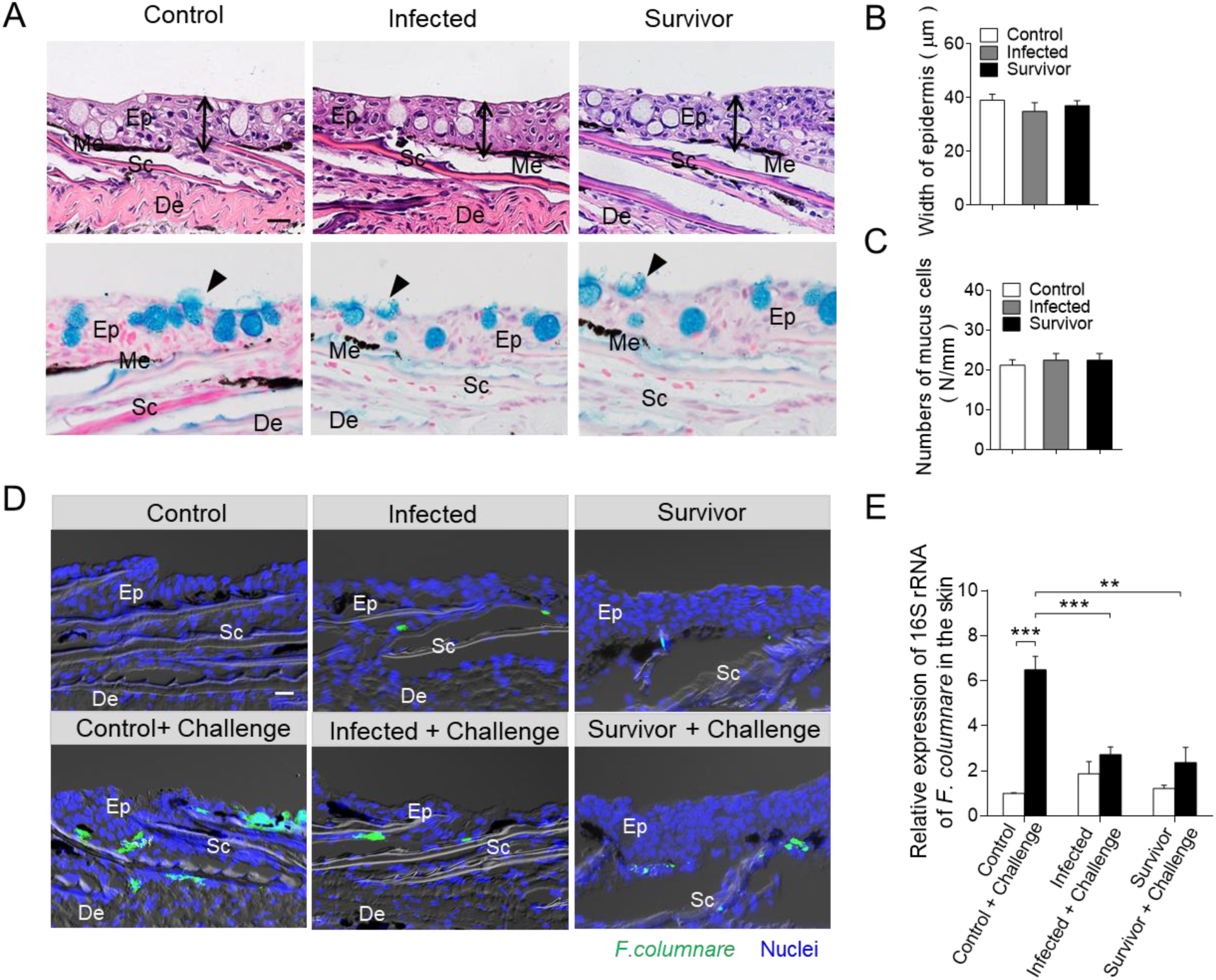
Bacterial load of the skin from survivor fish upon *F. columnare* re-infection. (**A**) Histological examination by H & E and AB staining of skin from fish infected with 28 days and survivor fish (*n* = 6). The black lines with double arrows and black triangle indicate the width of skin epidermis and the mucus secreting cells, respectively. Ep: epidermis, De: dermis, Sc: scale, Me: melanophores. Scale bars, 20 µm. (**B**) The width of skin epidermis of control fish, infected fish and survivor fish counted from A (upper) (*n* = 6). (**C**) The number of mucus cells per millimeter in the epidermis of trout skin from control, infected and survivor fish counted from A (lower) (*n* = 6). (**D**) DIC images of fluorescence analysis showed the GFP-*F. columnare* location and load in the skin from control and control-challenge fish (left), 28-day infected and 28-day infected-challenge fish (middle), survivor fish and survivor-challenge fish (right). These challenge fish were sampling at 4 days post re-infection with GFP-*F. columnare* (*n* = 6). Ep: epidermis, Sc: scale, De: dermis. Scale bars, 20 µm. (**E**) The relative expression of 16S RNA of *F. columnare* in the trout skin tissue of control, control-challenge, infected, infected-challenge, survivor and survivor-challenge group were measured by qRT-PCR (*n* = 6 fish per group). \**P* < 0.05, \*\**P* < 0.01, \*\*\**P* < 0.001 (unpaired Student’s *t*-test). Data are representative of at least three independent experiments (mean ± SEM).

### B cell responses and local proliferation in trout skin after bacterial infection

Our previous studies have demonstrated that IgT play a specialized role in skin mucosal immunity against an aquatic parasite (25). The decreases of *F. columnare*load in infected and survivor trout skin after re-challenge with a higher dose of the same pathogen, suggesting whether IgT play an important role in mucosal adaptive immune responses against this bacterial pathogen. To evaluate the role of skin mucosal Igs in response to bacterial challenge, we detected Ig and B-cell responses in trout skin infected with *F. columnare* by bath challenge. Using immunofluorescence analysis, we found very few IgT^+^ B cells as well as IgM^+^ B cells in the skin of control fish (Fig. 4A). However, fish infected after 28 days with *F. columnare*, a moderate (∼ 2-fold) but significant increase in the number of IgT^+^ B cells was detected in the skin (Fig. 4A, 4B). Importantly, when compared to control fish, we observed a notable accumulation of IgT^+^ B cells in the skin of survivor fish. In stark contrast to this, there was no change in the number of IgM^+^ B cells whether in infected or survivor fish groups when compared with that of control fish (Fig. 4A, 4B).

**FIGURE 4.**
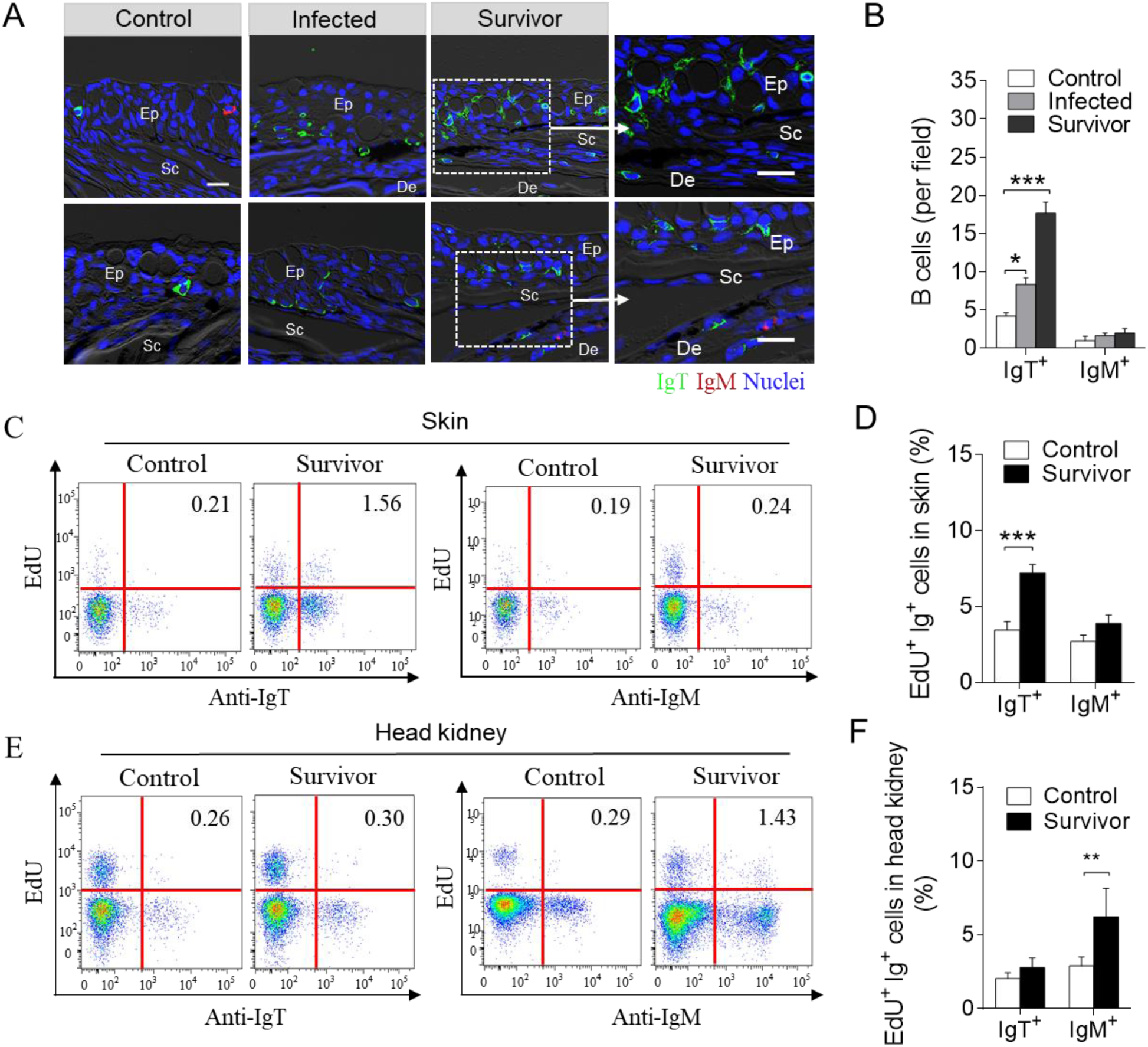
Accumulation and proliferation of IgT^+^ B cells in the skin of rainbow trout following *F. cloumnare* infection. (**A**) DIC images of immunofluorescence staining on trout skin paraffinic sections from control fish, infected and survivor fish stained for IgT (green) and IgM (red); nuclei are stained with DAPI (blue). Ep: epidermis, Sc: scale, De: dermis. Scale bars, 20 µm. (**B**) The number of IgT^+^ and IgM^+^ B cells in skin paraffin sections of control, infected and survivor fish infected with *F. columnare* are counted (*n* = 12, original magnification, ×20). \**P* < 0.05, \*\**P* < 0.01, \*\*\**P* < 0.001 (one-way ANOVA with Bonferroni correction). (**C**) Representative flow cytometry dot plot showing proliferation of IgT^+^ B cells and IgM^+^ B cells in skin leucocytes of control and survivor fish (*n* = 12). (**D**) Percentage of EdU^+^ cells from the total skin IgT^+^ or IgM^+^ B-cell populations in control and survivor fish (*n* = 12). (**E**) Representative flow cytometry dot plot showing proliferation of IgT^+^ B cells and IgM^+^ B cells in head kidney leucocytes of control and survivor fish. (**F**) Percentage of EdU^+^ cells from the total head kidney IgT^+^ or IgM^+^ B-cell populations in control or survivor fish (*n* = 12). \**P* < 0.05, \*\**P* < 0.01, and \*\*\**P* < 0.001 (unpaired Student’s *t*-test) in D and F. Data are representative of at least three independent experiments (mean ± SEM).

To confirm whether the substantial increase of IgT^+^ B cells in the survivor fish skin was due to proliferative IgT^+^ B cell in local skin mucosa or to the migration of these cells from other immune tissues such as systemic lymphoid organs and other MALT, the *in vivo*proliferation of B cells in control and survivor fish was detected by caudal vein injection of these fish with fluorescent EdU (5-ethynyl-20-deoxyuridine). Using flow analysis, the percentage of proliferating IgT^+^ B cells (EdU^+^ IgT^+^ B cells) showed a significant increase in the skin of survivor fish (∼ 7.20%) when compared to control fish (∼ 3.45%), while there was no change in the percentage of IgM^+^ B cells (EdU^+^ IgM^+^ B cells) between control and survivor fish (Fig. 4C, 4D). Interestingly, we found a notable rise of EdU^+^ IgM^+^ B cells in the head kidney of survivor fish (∼ 6.23%) compared to control fish (∼ 2.89%) while no differences in the proliferation of IgT^+^ B cells were detected (Fig. 4E, 4F).

### Bacteria-specific immunoglobulin responses in trout skin

To further study the skin Ig responses in trout skin after *F. columnare* infection, we performed SDS-PAGE and western blot analysis. Along with the significant increase of IgT^+^ B cells observed in the skin of infected and survivor fish, the IgT protein level in the skin mucus of the both groups were ∼ 8- and ∼ 15-fold higher respectively, when compared with that of control fish, while the levels of IgM and IgD did not differ (Fig. 5A). In serum, about 2-fold and an approximately 5-fold increases of IgT concentration were detected in infected and survivor fish, respectively. Conversely, the concentration of IgM in serum was much higher than that of IgT and had approximately 4- and 3-fold increases in infected and survivor fish, respectively, when compared with those of control fish (Fig. 5E). In contrast, the IgD protein concentration did not change significantly in either the skin mucus or serum of the same fish groups (Fig. 5A, 5E).

**FIGURE 5.**
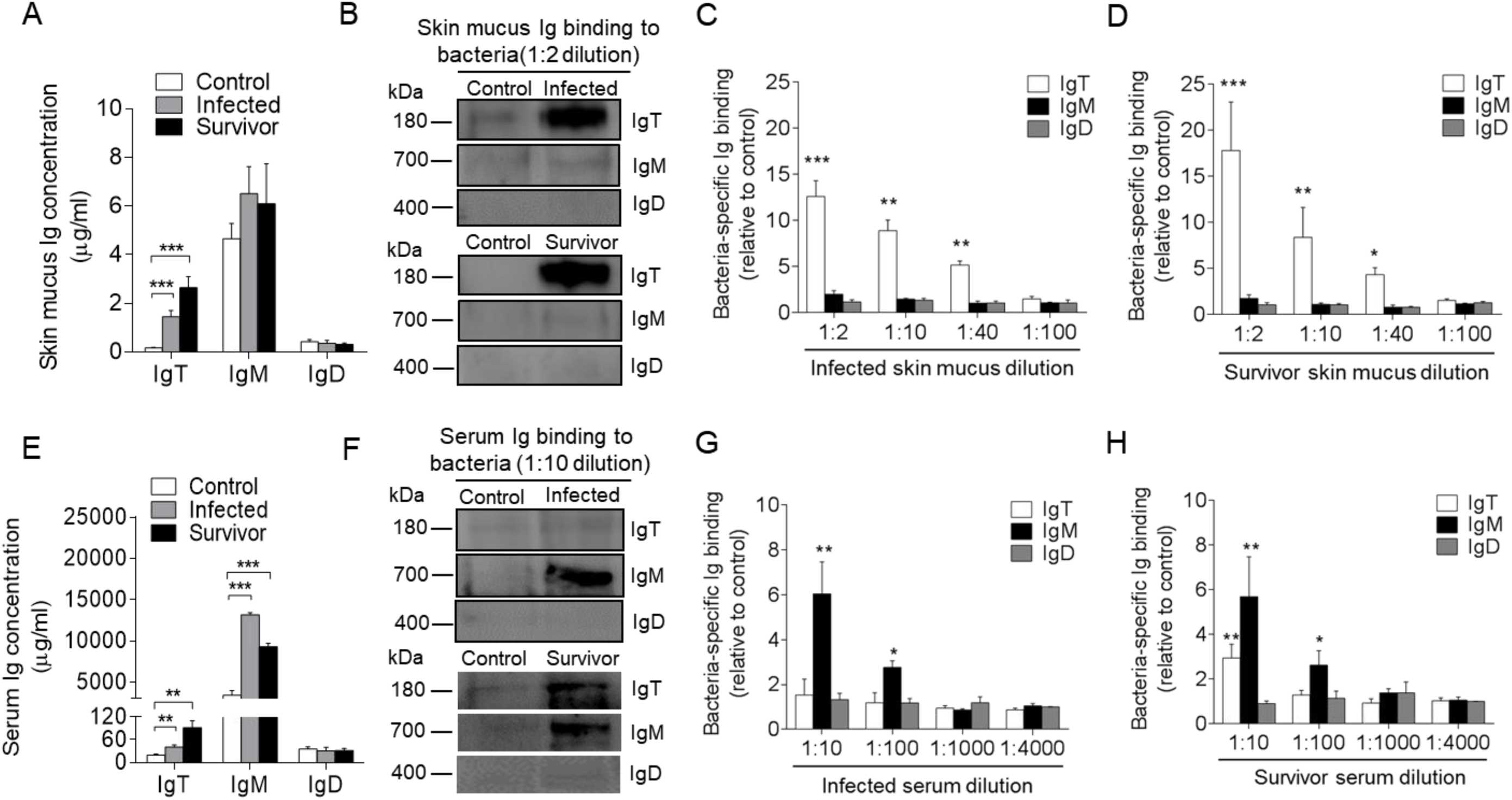
Ig responses in the skin mucus and serum from infected and survived fish.(**A**) Concentration of IgT, IgM, and IgD in skin mucus from control, infected, and survivor fish (*n* = 12). (**B**) Western blot analysis of IgT-, IgM- and IgD-specific binding to *F. columnare* in skin mucus (dilution 1:2) from infected and survivor fish. (**C** and **D**) IgT-, IgM- and IgD-specific binding to *F. columnare* in dilutions of skin mucus from infected (C) and survivor (D) fish evaluated by densitometric analysis of immunoblots and presented as relative values to those of control fish (*n* = 12-18). (**E**) Concentration of IgT, IgM, and IgD in serum from control, infected, and survivor fish (*n* = 12). (**F**) Western blot analysis of IgT-, IgM- and IgD- specific binding to *F. columnare* in serum (dilution 1:10) from infected and survivor fish. (**G** and **H**) IgT-, IgM- and IgD- specific binding to *F. columnare* in dilutions of serum from infected (G) and survivor (H) fish, evaluated by densitometric analysis of immunoblots and presented as relative values to those of control fish (*n* = 12-18). \**P* < 0.05, \*\**P* < 0.01, \*\*\**P* < 0.001 (unpaired Student’s *t*-test). Data are representative of at least three independent experiments (mean ± SEM).

The results of large increases in IgT^+^ B cells and IgT protein levels in the skin of infected and survivor fish suggested the generation of bacteria-specific IgT immune responses. To verify this hypothesis, using a pull-down assay, we measured the capacity of skin mucus Igs binding to *F. columnare*. Compared to control fish, we detected significantly higher (∼ 5-fold) bacteria-specific IgT binding up to 1/40 diluted skin mucus of infected and survivor fish, respectively (Fig. 5B–D). While bacteria-specific IgT binding was detected in serum with only in the 1/10 dilution of the survivor group (Fig. 5F, 5H). In contrast, ∼ 3-fold and ∼ 2.4-fold higher bacteria-specific IgM binding was observed in up to 1/100 serum dilutions both from infected and survivor fish, respectively (Fig. 5G, 5H). However, bacteria-specific IgD binding is hardly detected in either the skin mucus or the serum from infected and survivor fish (Fig. 5A–H).

To test further the local production of pathogen-specific Igs in trout mucosal and systemic tissues respectively, we analyzed bacteria-specific Ig titers of culture medium derived from cultured skin, head kidney, and spleen explants of control and survivor fish (Fig. 6A–6F). We detected bacteria-specific IgT binding up to the 1/100 dilution of medium of cultured skin explants from survivor fish, while very low bacteria-specific IgM responses (Fig. 6A, 6D) were detected (only in the 1/2 dilution in the same medium). In contrast, dominant bacteria-specific IgM binding (up to the 1/10 dilution) was observed in the medium of survivor head kidney and spleen explants, while similar (up to the 1/10 dilution) and lower (up to the 1/2 dilution) bacteria-specific IgT responses were detected in the medium of survivor head kidney and spleen explants respectively (Fig. 6B, 6E, 6C, 6F). Interestingly, we could not detect any bacteria-specific IgD binding in either the mucosal (skin) or systemic (head kidney and spleen) tissue explants from survivor fish (Fig. 6A–6F).

**FIGURE 6.**
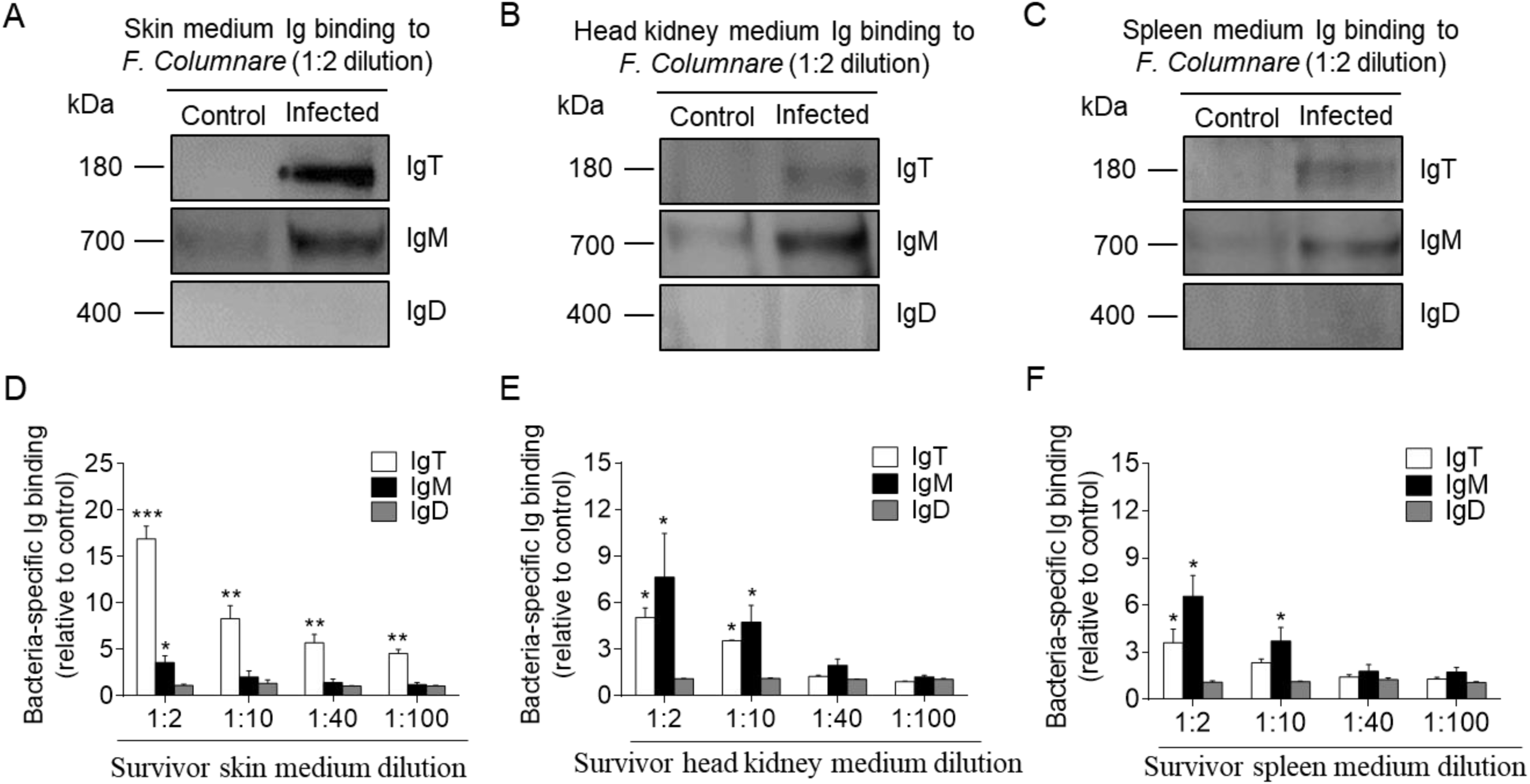
Local IgT-, IgM- and IgD-specific responses in skin explants of survivor fish.(**A**–**C**) Skin, head kidney, and spleen explants (∼ 50 mg each) from control and survivor fish were cultured in medium (400 μl) for 7 days. Immunoblot analysis of IgT-, IgM- and IgD- specific binding to *F. cloumnare* in the culture medium of skin (A), head kidney (B), and spleen (C) (dilution 1:2) from control and survivor fish. (**D**–**F**) IgT-, IgM- and IgD- specific binding to *F. cloumnare* in dilutions of culture medium from skin (D), head kidney (E), and spleen (F) from control and survivor fish, measured by densitometric analysis of immunoblots and presented as relative values to those of control fish (*n* = 12). \**P* < 0.05, \*\**P* < 0.01, and \*\*\**P* < 0.001 (unpaired Student’s *t*-test). Data are representative of at least three independent experiments (mean ± SEM).

### pIgR in trout skin after bacterial infection

Previous studies have indicated that rainbow trout pIgR may take part in the skin mucosal immune responses to parasitic infection by mediating the transepithelial transport of secretory Ig (9). Thus, we hypothesized that pIgR might also mediate the transport of IgT to the skin mucus during immune responses to *F. columnare*. Here, using immunofluorescence analysis, we observed that some of the skin epithelial cells from control fish stained by the anti-trout pIgR antibody (Supplemental Fig. 3A). Importantly, the number of pIgR^+^ cells significantly increased in the epidermis of infected (∼ 1.6-fold) and survivor fish (∼ 2.4-fold) (Supplemental Fig. 3A, 3B) when compared with those of control fish. Using qRT-PCR, the expression of pIgR was found to be significantly upregulated in the skin of infected (∼ 3-fold) and survivor fish (∼ 6.6-fold), respectively, when compared with that of control fish (Supplemental Fig. 3C). Moreover, the concentration of pIgR in skin mucus post bacterial infection were also detected and calculated by western blot analysis. Our results showed that the relative fold changes of pIgR in infected and survivor fish when compared to that of control fish, were significantly increased (∼ 1.7-fold for infected group and ∼ 3.4-fold for survivor group; Supplemental Fig. 3D, 3E).

## Discussion

Fish skin, is the outermost organ of the organism’s body, which is continuously exposed to a wide array of potentially harmful insults in the aquatic environment. This constant exposure to microbes has led to the evolution of a mucosal immune network in the teleost SALT, which ensures that adequate immune responses are mounted upon antigenic challenge while maintaining overall tissue and microbiota homeostasis (16, 40–42). However, till now little is known with regards to the evolution of skin mucosal B cell and antibody responses to bacterial pathogens in early vertebrates. Most of the literature in that area for teleost fish predates the discovery of IgT as the key mucosal immunoglobulin in these species (10). In that regard, some studies in several teleosts have reported low levels of pathogen-specific IgM titers upon bacterial infection (43, 44). Moreover, whether the skin mucosa behaves as an inductive or effector site of mucosal Ig responses has never been investigated. Here, we report for the first time in a teleost that infection with *F. columnare*, a widespread fish bacterial pathogen, can elicit local skin proliferative IgT^+^ B cell responses as well as significant bacterial-specific mucosal IgT titers.

Columnaris disease, caused by the Gram-negative bacterium *F. columnare*, severely impacts the global production of many fish species (33, 45). As a mucosal bacterial pathogen, *F. columnare* infection results in skin epithelial erosion, gill necrosis and ulcers, with a high degree of mortality (33, 45). To evaluate mucosal immune responses to *F. columnare*, we exposed fish to this pathogen following a well-established waterborne challenge model with a novel fluorescent (GFP) strain of *F. columnare* (37). Importantly, GFP-*F. columnare* exhibited similar growth and virulence properties as the wild-type bacteria, and thus, this newly constructed GFP-*F. columnare* represents a new reagent that is expected to significantly advance the study of host- *F. columnare* interactions. At 4 dpi, clinical signs were detected in the bacteria-infected rainbow trout, and qRT-PCR and immunofluorescence further proved the successful invasion of GFP-*F. columnare* into rainbow trout. It is worth point that the use of our unique GFP-*F. columnare* strain was extremely instrumental in assessing the capacity of this pathogen to intrude into the skin epidermis of fish under the experimental conditions tested. Importantly, upon infection, we observed significantly decreased numbers of mucus cells and thickness of the epidermis in the trout skin. Thus, our results suggest that *F. columnare*’s successful invasion relies on destroying the epidermal layer structure of fish skin, which is in line with what has been shown in previous studies (47, 49). However, the mechanisms of bacterial infection still need to be further investigated. In agreement with the noteworthy histopathological changes in trout skin, we found that 15 immune-related genes were significantly upregulated in trout skin as early as 1 dpi when compared with those in the head kidney and spleen, which indicates that the immune responses in trout skin are activated significantly earlier than in systemic lymphoid tissues, thus supporting further the notion of the skin acting as the first line of defense against bacterial infections in fish. Importantly, both innate and adaptive immune molecules were induced in the skin upon *F. columnare* infection, including antibacterial peptides, cytokines, chemokines, complement factors, and Igs. Thus, our results provide a general picture of the predominant immune responses that take place in the trout skin after *F. columnare* infection. Of these responses, it is worth pointing the increased transcript levels of cathelicidin 1 and proinflammatory cytokines such as IL8 and TNFα found in trout skin immediately after *F. columnare* infection. Moreover, the anti-inflammatory cytokine IL10 was also induced after bacterial infection. Combined with previous studies (50–53), our results suggest that *F. columnare* infection induces a significant inflammatory reaction as well as strong immune responses in the fish skin epidermis early upon infection. In contrast, these types of responses occur in the dermis of mammals (16, 54), probably reflecting the different physiological needs of fish and mammalian skin tissues with regards to the water or terrestrial environments where these species live.

The use of GFP-*F. columnare* enabled us to find that the skin epithelium of trout previously exposed and thus, immune to *F. columnare*, cannot get intruded upon re-exposure to this pathogen. These findings suggested that local humoral immunity might be elicited in trout SALT thus contributing to protection upon re-infection. Here we first showed significant increases in the concentration of IgT but not IgM in the skin mucus of infected and survivor fish exposed to *F. columnare*. These data correlated with the large accumulation of IgT^+^ but not IgM^+^ B cells in the skin epidermis of the same animals. These results paralleled those obtained in a previous study where fish surviving from a parasite infection exhibited large accumulations of IgT and IgT^+^ B cells in the skin epidermis (9). It is worth mentioning that upon bacterial infection, several mammalian MALTs have been shown to have similar significant accumulations of IgA^+^ B cells (55–59). In humans, several studies have described a correlation between resistance to *Vibrio cholerae* infection and high specific sIgA titers (60, 61). Importantly, our findings show for the first time in fish the detection of bacteria-specific IgT titers mainly in the skin mucus and to a much lesser degree in the serum of survivor fish. Although the kinetics of IgM production in the skin following bacterial infection had been investigated in some species (62–65), till now the role of sIgT in skin bacterial infections had not been addressed, probably due to the lack of specific antibody reagents recognizing IgT (66, 67). Moreover, we hardly found any bacteria-specific IgM titers in skin mucus of infected and survivor fish, which were instead detected almost exclusively in the serum in very high levels. Therefore, we can preliminarily conclude that skin IgT and IgT^+^ B cell responses against *F. columnare* are specifically confined in its mucosa, whereas IgM responses are overwhelmingly found in the serum, and thus of systemic nature. Interestingly, previous studies in trout with a similar bacterial pathogen (*Flavobacterium psychrophilum*) have shown that bath exposure of trout to this pathogen could not elicit bacteria-specific IgM responses in the trout skin mucus (68), as we have also found for *F. columnare*. Thus, the authors speculated that perhaps IgT is the immunoglobulin involved in skin responses against *F. pychrophilum*, a hypothesis that seems to be supported by our data, although that hypothesis remains to be investigated.

The accumulation of IgT^+^ B cells and high bacteria-specific IgT titers observed in the skin mucosa from infected and survivor fish led us to hypothesize the local proliferation and induction of IgT^+^ B cells and IgT antibody responses respectively upon bacterial infection. Confirming this hypothesis, we found strong local proliferative IgT^+^ B cells responses in the teleost skin but not in the head kidney of the same fish, suggesting that the accumulation of IgT^+^ B cells in their skin is due to local proliferation rather than migration of these cells from the systemic immune organs. We cannot rule out however the possibility that these IgT^+^ B cells are induced in other mucosal surfaces, and upon migration to the SALT, they start proliferating. Future work will have to address the aforementioned possibility. Critically, the production of significant titers of bacteria-specific IgT in skin explant cultures confirmed that these IgT antibodies are produced locally and suggest the presence of specific plasma cells in the local skin mucosa. Overall, considering that *F. columnare* firstly invades and stays in the skin epidermis, it is unlikely that the inductive site is a systemic lymphoid tissue, thus our results imply that trout SALT is an inductive site of the observed IgT mucosal immune responses. We have previously shown similar results in the mucosa of gill and nose when exposed to pathogenic challenge (7, 8). It is worth noting also that similar results have been shown in mucosal inductive sites of some mammalian MALTS, including the GALT, NALT and bronchus-associated lymphoid tissue (BALT) where a local mucosal response governed by a mucosal Ig (sIgA) was strongly induced upon bacterial infection, and in contrast, the systemic response is dominated by IgG/IgM (69–71).

A key feature of pIgR is the mediation of the transepithelial transport of secretory Igs into the mucosal surfaces in both mammals and teleost (2, 72). Previous studies have shown that the putative trout secretory component (tSC) of pIgR is associated with mucosal IgT and IgM in gut (10), skin (9), gills (7), and nasal mucus (8), contributing to the Ig response to parasite infection. Here, we found that trout pIgR (tpIgR) was mainly expressed in the epithelial layer of the skin from control, infected, and survivor fish. Critically, here we see a significant increase in the numbers of pIgR^+^ cells and transcript levels of pIgR in the skin from bacterial infected trout when compared with that of control fish. Interestingly, similar results have been reported in mammals, for example, the expression of pIgR was found upregulated in germ-free mice implanted with *Bacteroides thetaiotaomicron* (73). In addition, pIgR expression can be upregulated in HT-29 cells by reovirus (double-stranded RNA) (74) and bacteria (*Enterobacteriaceae*) with LPS on the surface (75). Thus, our results indicate such increases in the levels of pIgR upon pathogenic challenge is an evolutionary conserved feature from fish to mammals which probably has the role to increase the transport rate of mucosal immunoglobulins to mucosal surfaces upon pathogenic insult.

In conclusion, our study provides the first evidence of local B cell proliferation and IgT secretion in teleost skin, which suggests that teleost SALT is an inductive site of mucosal B cell and immunoglobulin responses. Based on our studies, we propose a model in which upon bacterial infection, epidermis damage occur which leads to an up regulation of immune-related genes in the skin as well as to processes of IgT^+^ B cell activation and proliferation within the SALT, resulting into the production of bacteria-specific IgT (Fig. 7). Thus, from an evolutionary viewpoint, our results reinforce the formerly stated view (76) that regardless of their phylogenetic origin and tissue localization, specialized mucosal Igs (i.e., IgA, IgT) of all vertebrate MALT-containing surfaces function under the guidance of primordially conserved principles. Moreover, our findings in this phylogenetically primitive vertebrate strongly suggest a universal requirement for vertebrate MALT-containing mucosal surfaces to use a dedicated Ig for the maintenance of homeostasis. Additionally, since many fish pathogens enter the host through the skin mucosa, the knowledge derived from our findings also has special relevance from a more practical perspective, as it may lead to the development of fish vaccines that can effectively induce bacteria-specific IgT immune responses in skin.

**FIGURE 7.**
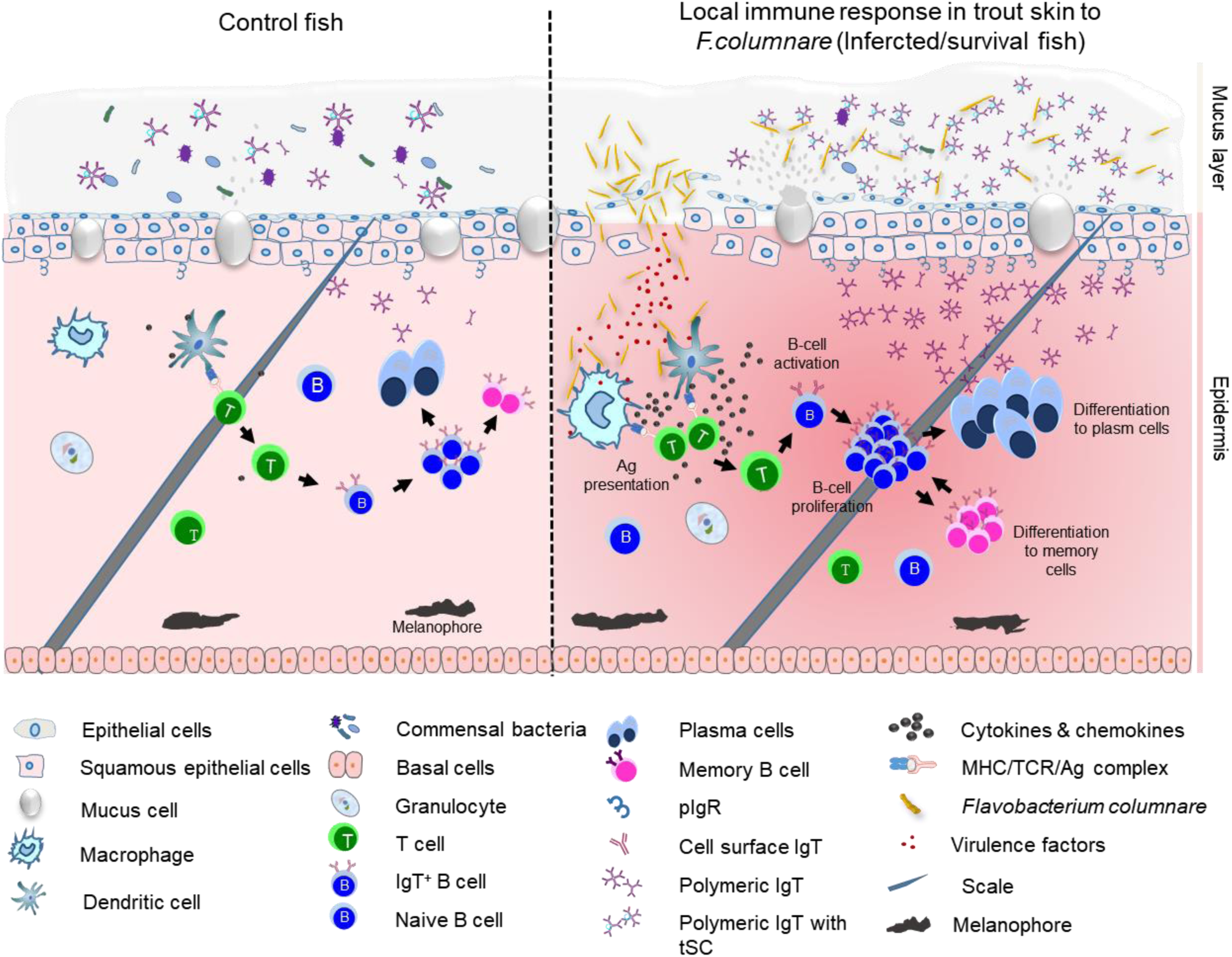
Proposed model of local IgT and IgT^+^ B cell induction in the skin after *F. columnare* infection. The presumed structure of skin displaying mucus layer, epithelia layer and dermis. The proposed model contains two partitions: control (left) and infected/survivor (right). Induction of local IgT responses in the trout skin based on our findings. On the left is the scheme of a typical skin structure in control (naive) teleost fish. The number of IgT^+^ B cells in control fish skin is low. IgT are produced by IgT–secreting B cells and transported from the epithelium into the skin mucus layer via pIgR. The secreted IgT coats the majority of microbiota in the skin surface. In the skin epidermis, abundant mucus cells are present, and their production contains antibacterial molecules to protect host against pathogens. Upon *F. columnare* invasion, their antigen (Ag) can be taken up by antigen-presenting cells (APC) and presented to naive CD4^+^ T cells. Then IgT^+^ B cells are activated by Ag-specific CD4^+^ T cells and start to proliferate locally and differentiate into plasma cells (PCs) in skin epithelial layer. These differentiated IgT–secreting cells can produce large amounts of pathogen-specific IgT, which can be transported by pIgR into skin mucus layer where those pathogen-specific IgT can specifically binding to the bacteria *F. columnare*. Alternatively, some IgT^+^ plasma cells may differentiate further into memory IgT^+^ B cells. When *F. columnare* invade the host again, the memory IgT^+^ B cells would directly proliferate and differentiate into plasma cells, and then rapid produce specific–IgT to binding *F. columnare*.

## Acknowledgements

We thank Dr. J. Oriol Sunyer (University of Pennsylvania) for his generous gift of anti-trout IgM, anti-trout IgD, anti-trout IgT mAbs, anti-trout IgT and anti-trout pIgR pAbs.

## Supplemental information

**Supplemental Fig 1.**
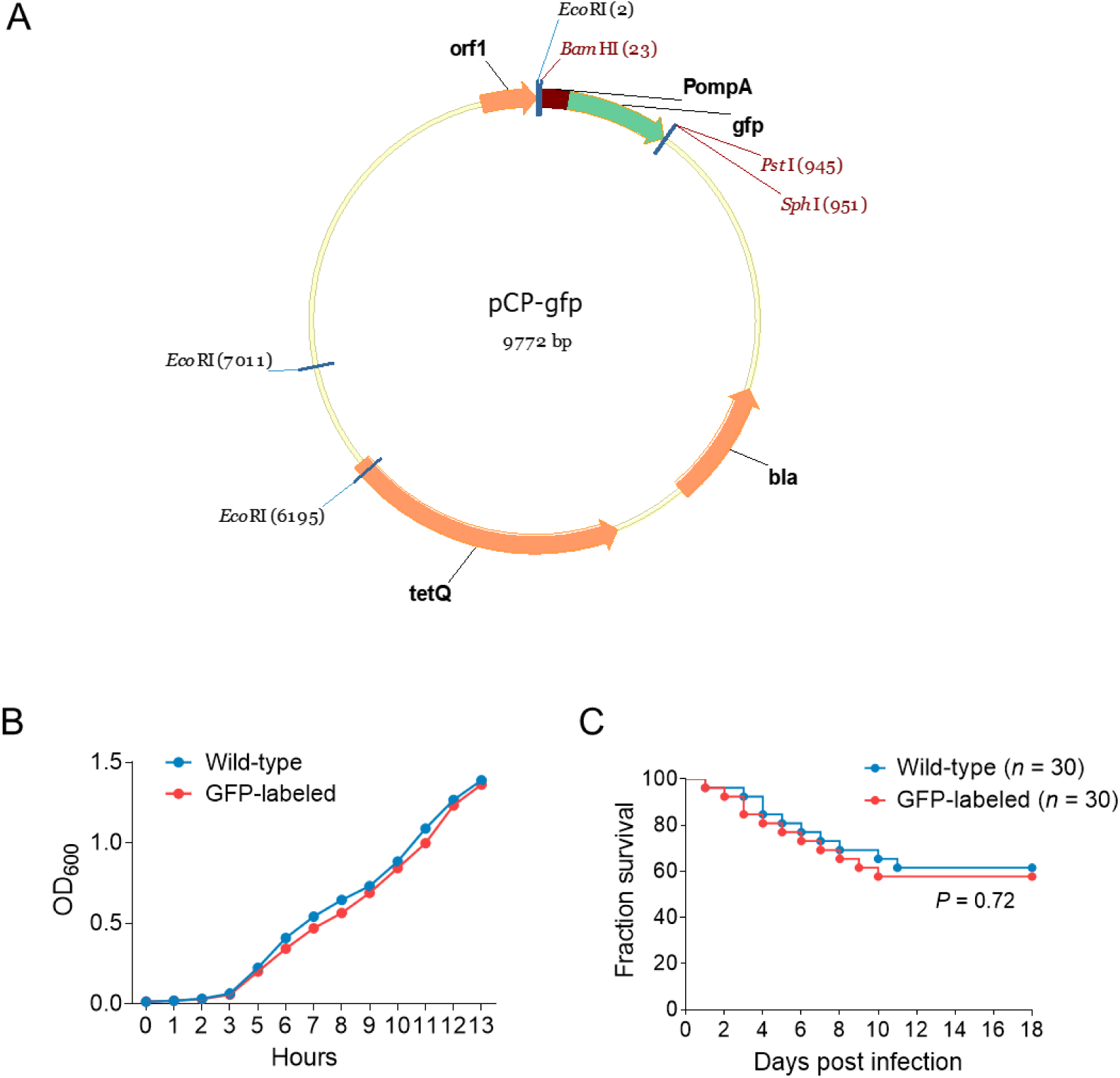
Construction of GFP-*F. columnare* G_4_ strain. (A) Map of the plasmid pCP23-gfp. (B) Growth curves for wild-type *F. columnare* and the GFP-*F. columnare*. (C) Survival curves of fish infected with wild-type G_4_ and the GFP-*F. columnare* (*n* = 30). Data are representative of at least three independent experiments (mean ± SEM).

**Supplemental Fig 2.**
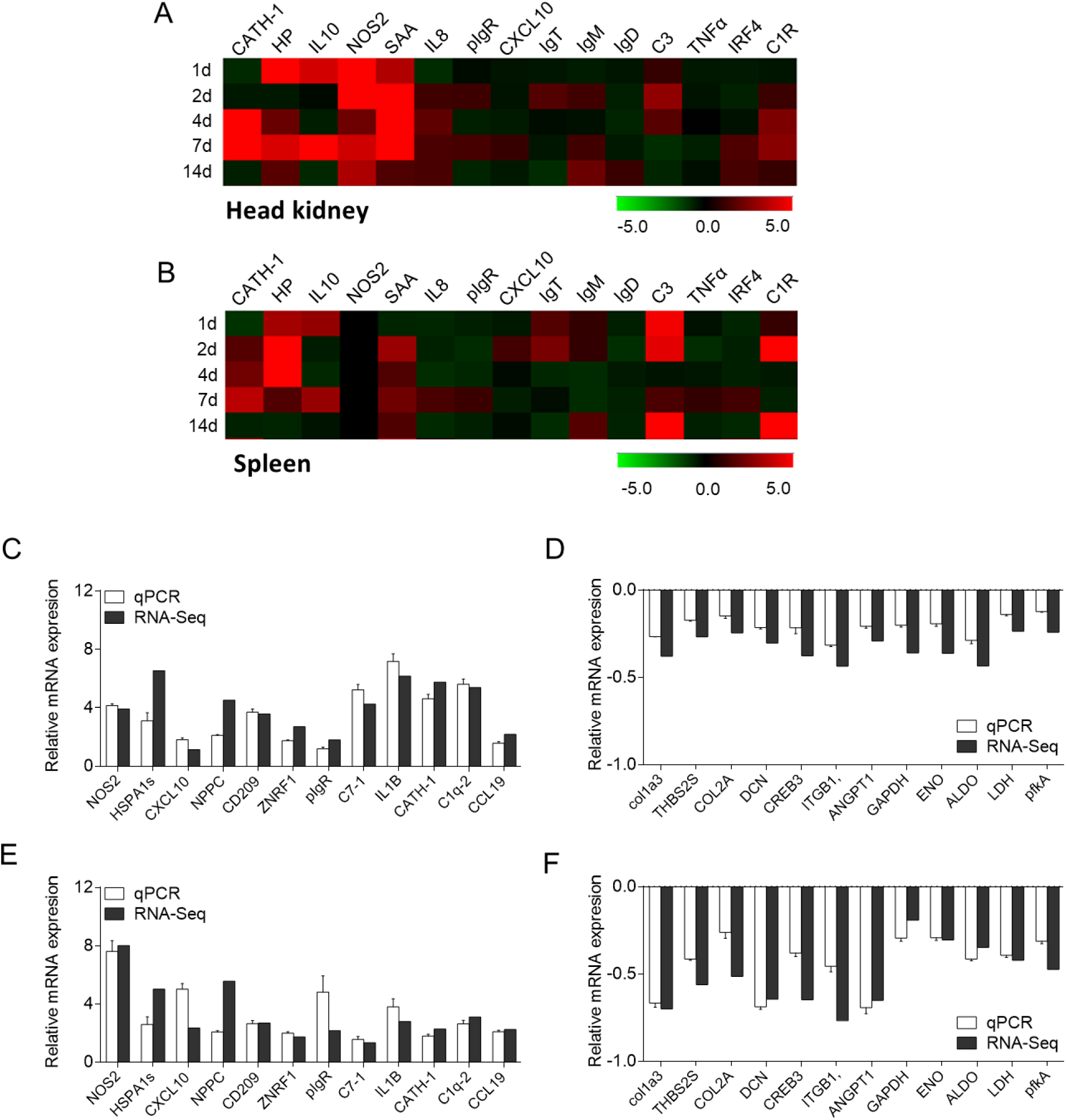
Immune responses in trout spleen and head kidney following *F. columnare* infection, and validation of transcriptomic data by PCR (A and B) Heat map illustrates results from qRT-PCR of mRNAs for selected immune markers in *F. columnare*-infected fish versus control fish measured at 1, 2, 4, 7, and 14 dpi in trout head kidney (A) and spleen (B) (*n* = 6). Data are representative of three different independent experiments (mean ± SEM). (C–F) Transcriptomic differential expressed genes in experimental groups at 2 (C and D) and 14 days (E and F) post infection were randomly chosen and detected using qPCR to validate RNA-seq. Positive numbers in the Y axis mean up-regulated, while negative values mean down-regulated (*n* = 6). Data are representative of three different independent experiments (mean ± SEM).

**Supplemental Fig 3.**
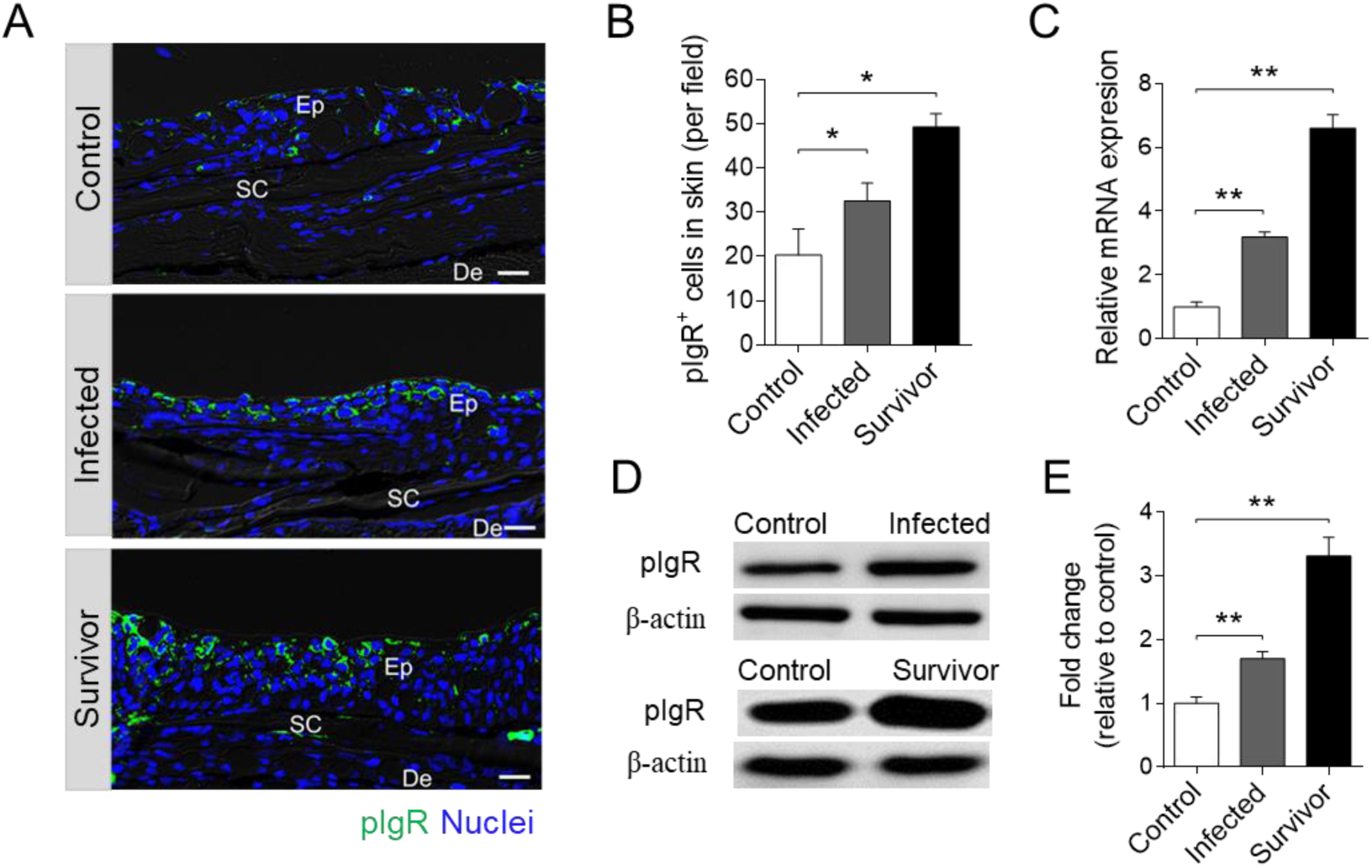
Accumulation of pIgR and pIgR^+^ cells in the skin of rainbow trout infected with *F. columnare*. (**A**) DIC images of immunofluorescence staining on trout skin paraffinic sections from control fish, infected, and survival fish, stained for pIgR (green) and nuclei with DAPI (blue) (*n* = 12). Ep: epidermis, De: dermis, Sc: scale. Scale bars, 20 µm. (**B**) The number of pIgR^+^ cells in skin paraffin sections of control, infected, and survived fish are counted (*n* = 12, original magnification, ×20). (**C**) Relative mRNA expression of pIgR in the skin of control fish, infected, and survival fish were detected by qRT-PCR (*n* = 6). (**D**) Immunoblot analysis of pIgR concentration in skin mucus from control, infected and survivor fish following *F. columnare* infection. (**E**) Relative concentration of pIgR from infected and survivor trout skin mucus to that of control fish (*n* = 12). \**P* < 0.05, \*\**P* < 0.01 (unpaired Student’s *t*-test). Data are representative of at least three independent experiments (mean ± SEM).

**Supplemental Table I.**
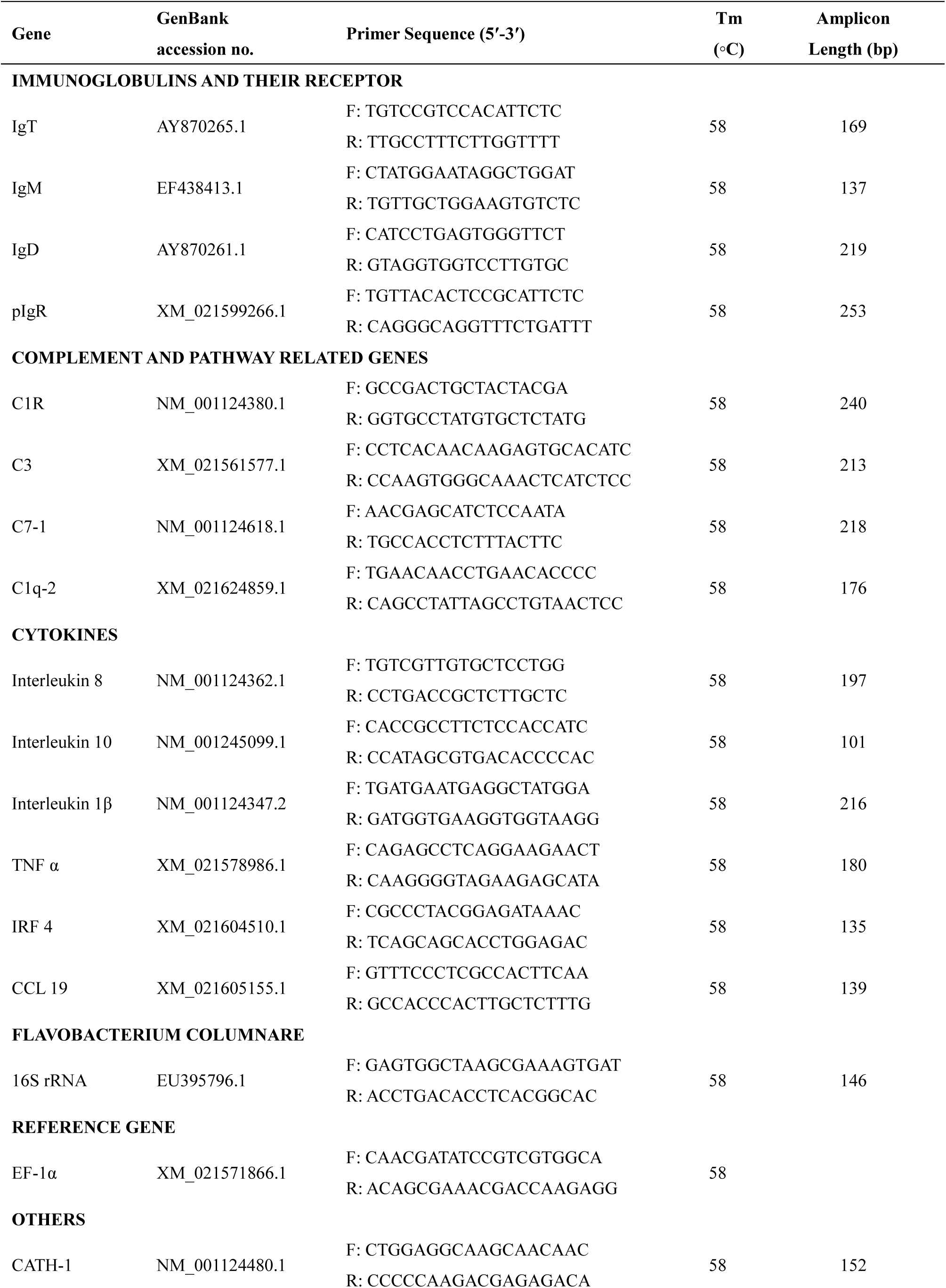
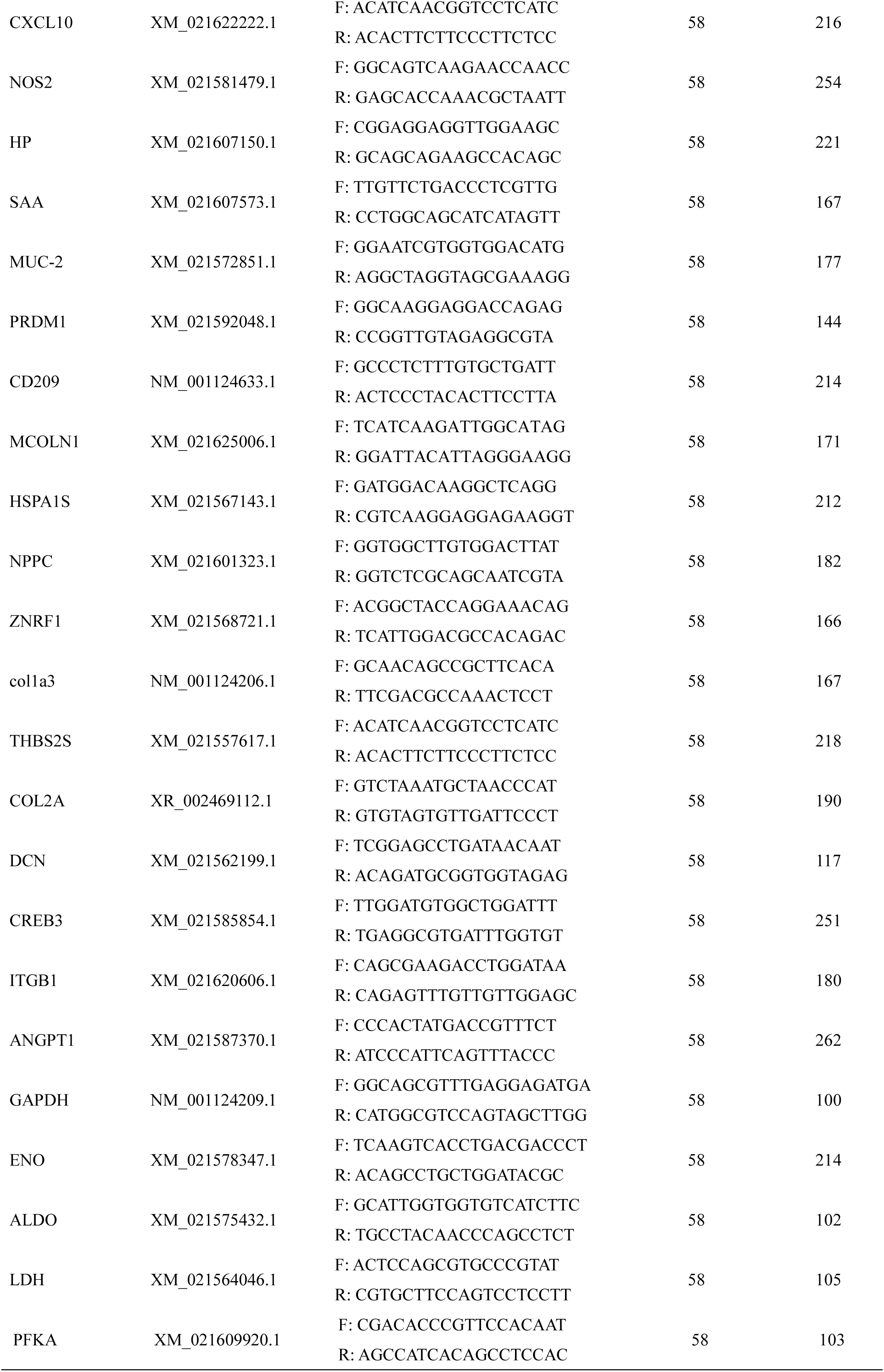
List of primers for real-time quantitative PCR amplifications.

